# Development and characterization of a miniaturized focused ultrasound extraction (FUSE) system to enable rapid, point-of-care genetic testing

**DOI:** 10.64898/2026.09.24.754152

**Authors:** Alexia Stettinius, Adam Maxwell, Kayleigh Canavan, MoonWon Jeon, Daniel Yang, Gabby Lopez, Annika Griggs, Misa Winters, Qian Zhang, Jason Holliday, Eli Vlaisavljevich, Hal Holmes

## Abstract

Point-of-care (POC) molecular detection has enabled rapid and accessible testing at or near the site of need, facilitating timely decision-making in clinical, field, and industrial settings. Despite these advancements, molecular detection platforms remain limited in their ability to prepare DNA from complex tissue matrices, restricting their applicability to a narrow range of sample types. To address sample preparation inefficiencies, focused ultrasound extraction (FUSE) has been developed for the rapid release of DNA from robust tissue matrices. FUSE disintegrates target tissue and releases DNA by delivering high pressure ultrasound pulses that generate a cavitation bubble cloud. In prior work, FUSE was performed using large, laboratory-based equipment. Here, a miniaturized FUSE device was designed and tested to enable POC DNA sample preparation. Design constraints were defined based on prior FUSE experiments, and based on these metrics, a 750 kHz cylindrical transducer was fabricated. The pressure output was measured, and high-speed optical imaging demonstrated that the transducer generated sustained cavitation within the defined focal region. The feasibility of DNA extraction from piscine and timber tissues was evaluated, and results show that the device can release DNA with quality suitable for PCR-based detection from both tissue types. These results demonstrate the potential of this device to rapidly extract DNA from complex samples and expand the applicability of molecular detection platforms.

## 1. Introduction

The rapid advancement and widespread adoption of nucleic acid amplification tests (NAATs) have transformed the landscape of molecular diagnostics, particularly for point-of-care (POC) applications [1]. NAATs encompass a range of amplification methods, including polymerase chain reaction (PCR), loop-mediated isothermal amplification (LAMP), and recombinase polymerase reaction (RPA), among others [1, 2]. Together, these technologies enable sensitive and specific molecular detection across a broad range of applications, including healthcare, forensics, biosecurity, and agriculture [3–7]. Advances in assay speed, power requirements, and platform scalability have further expanded the use of NAATs beyond centralized laboratory settings and toward POC testing [1, 8]. This portability enables molecular testing across diverse environments, including at-home use, major hospitals, laboratories, mobile healthcare clinics, or remote field settings for forensic, agricultural, and environmental security use cases [9–12].

Despite the advantages of NAATs, DNA sample preparation shortcomings restrict the full potential of these platforms. Although some NAAT platforms incorporate integrated sample processing and DNA extraction [13, 14], effective preparation of complex samples remains a major challenge, particularly for robust tissues and samples rich in carbohydrates, metabolites, and other amplification inhibitors. Examples include plants, fibrous tissue, formalin-fixed paraffin-embedded (FFPE) tissue, Gram-positive bacteria, and processed tissues [15–19]. Current DNA sample preparation methods for these sample types have substantial time and resource requirements, often requiring a fully equipped laboratory and more than 24 hours for DNA extraction [20–22]. Incorporating the physical processing required for these complex samples into NAAT platforms therefore remains challenging while preserving the portability, small footprint, and rapid turnaround required for POC testing.

To overcome DNA extraction shortcomings and enable the development of NAATs integrated with robust sample preparation workflows, focused ultrasound extraction (FUSE) has been developed as a novel method for sample processing and DNA release. FUSE utilizes short-duration, high amplitude focused ultrasound pulses to generate a cavitation bubble cloud capable of breaking down samples on the tissue and cellular level for tissue homogenization and DNA release. Foundational studies have validated the feasibility of FUSE for preparing DNA from biological tissues and plant tissues (leaves, timber), representing samples with a range of complexity and applications [23–26]. Further, FUSE enables rapid sample processing, with the release of high-quality DNA extracts in as little as ten seconds. The time efficiency of FUSE, combined with its ability to simultaneously homogenize tissue and lyse cells without mechanical components or heated incubation, highlights its potential for integration with NAAT platforms.

However, prior FUSE studies were performed using a 32-element 500 kHz array transducer that was originally designed to perform histotripsy for non-invasive medical procedures [27, 28]. The large aperture size of this transducer (∼12 cm) limited its portability and required the transducer to be suspended in a large water tank with samples positioned in the focus 7.5 cm from the face of the transducer. This setup ultimately required the use of sizeable, complex, laboratory-based equipment, demonstrating the need to scale down the FUSE experimental configuration to enable portable sample processing. There have not been any studies to date investigating the feasibility of FUSE DNA sample preparation with alternate transducer designs and acoustic parameters.

In this study, we developed, fabricated, and tested a miniaturized FUSE system to broaden the accessibility of complex DNA sample preparation. First, the device was designed around key user and acoustic requirements to enable consistent and controlled delivery of acoustic energy to the sample. The transducer output was then simulated to assess whether the resulting acoustic pressure field could achieve sufficient focal gain to sustain cavitation. The transducer was fabricated and characterized using electrical, acoustic, and high-speed imaging techniques to evaluate its electrical impedance, focal pressure field, and cavitation cloud dimensions. The device was tested using piscine and timber samples, and performance was evaluated based on DNA yield, purity, and qPCR success and efficiency. We hypothesized that a miniaturized FUSE system could generate consistent cavitation and achieve effective DNA extraction from complex tissues, with quantities and quality sufficient for qPCR-based detection.

## 2. Materials and Methods

### 2.1 Device Design Specifications

A first-generation miniaturized FUSE prototype was designed to ensure robust tissue processing and efficient DNA release, adhering to the following criteria:

1. sufficient focal gain to generate FUSE cavitation clouds;
2. a focal pressure zone limited to the internal dimensions of a small test tube;
3. compatibility with a sample holder with simple alignment, supporting a test tube in the acoustic focus;
4. compactness and durability for easy portability.

The primary design objective was to develop a small transducer geometry capable of generating acoustic cavitation in a sample tube. Based on prior studies, it was assumed that the sample tube would have an inner diameter between 6.35 and 9.5 mm. The height, defined as the tube height occupied by the sample and lysis buffer, was dependent on the sample and lysis buffer volume. The height was expected to range from 11 to 16 mm, varying based on sample type and application [23, 25, 26]. For the generation of sustained FUSE cavitation clouds and to isolate cavitation to the test tube, it is necessary to restrict the focal zone (-6 dB) beamwidth to be similar to or smaller than the internal tube diameter and height [29–32]. To meet these constraints, the device was designed to limit the cavitation cloud dimensions to 11 mm along the major axis and 6 mm along the minor axis. Additional design considerations included a sample holder that preserved unobstructed acoustic window for energy delivery to the sample, as well as minimizing the transducer focal length to reduce the overall device footprint.

### 2.2 Device Simulation and Fabrication

Based on the design constraints, a single element cylindrical transducer with a concentric focusing lens was simulated using a 1D piezoelectric element model (KLM model) [33] and the k-Wave open source toolbox [34], a k-space pseudospectral time domain method to model the acoustic output. This design enabled spherical focusing from a cylindrical source. The KLM model specified a 400 V_pp_ square-wave signal with a frequency of 750 kHz was delivered to an L-bridge matching network with capacitance and inductance of 6000 pF and 7 μH, respectively, and then propagated to the transducer [35]. The mechanical impedance of the transducer was determined by assuming a lens with an inner radius of 19.25 mm and thickness of 2.9 mm (PerFORM, Ceramic-Like Advanced HighTemp, Protolabs, Maple Plain, MN, USA) and was concentrically aligned with the piezoelectric element with a 3 mm thickness (Steiner and Martins, Davenport, FL, USA). This model was used to simulate source pressure output of the transducer.

The acoustic pressure field was then simulated using the k-Wave open source toolbox. The acoustic source was modeled as an axisymmetric element with a 22.5 mm radius of curvature along the height of the cylinder with a cylinder height of 14 mm. The delivery of a 40-cycle pulse with a source pressure *p_0_* = 0.2 MPa through a coupling medium (water) was simulated. The coupling medium was bounded by air on the top surface and outer diameter of the cylinder, with a thin layer of plastic at its bottom base to contain the coupling medium in the internal cavity of the cylinder. The pressure output in the acoustic field was simulated over 200 μs, and the temporal and spatial location of the peak negative pressure (*p-*) was determined. This simulation was performed in an axisymmetric domain with the axis of symmetry equivalent to the central axis of the cylinder. In this way, the entire cross-section of the acoustic focal zone was visualized.

A solid model of the device was developed based on simulation results. This included a focusing lens with a closed bottom placed within the cylindrical source to create an exposure chamber. Sample holders were designed to support acoustically permeable tubes of varying sizes and fit concentrically within the chamber. Lastly, external housing was developed to contain the transducer and sample holder. A solid rendering of the FUSE device was constructed using CAD models (Autodesk Inventor, San Francisco, CA, USA) (**Figure 1**).

**Figure 1.**
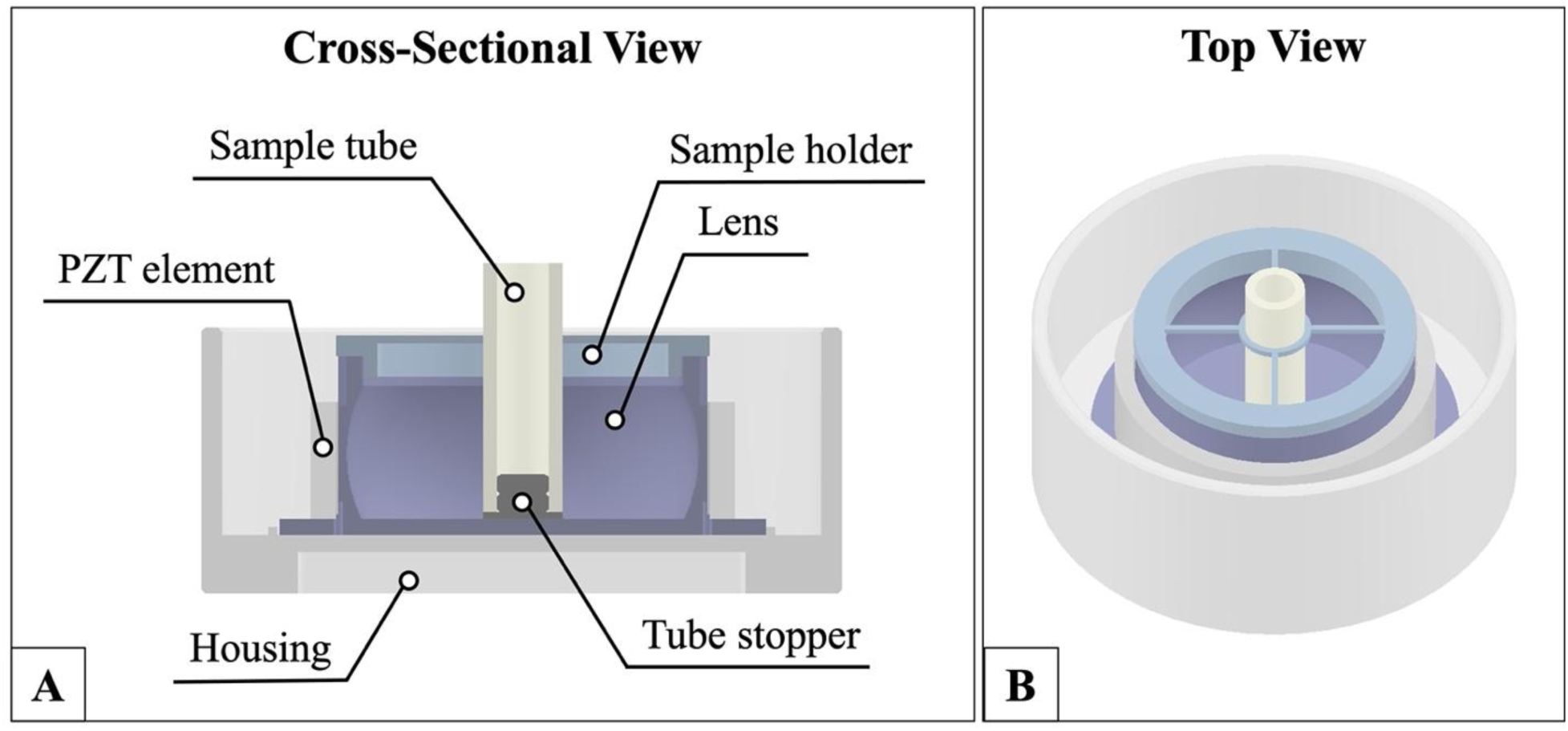
FUSE device design. (A) Cross-sectional and (B) solid CAD models of the device showing the transducer, sample holder apparatus, and protective housing.

The transducer was fabricated using a piezoceramic cylinder element with an inner diameter of 45 mm and a height of 14 mm (Steiner and Martins, Davenport, FL, USA). The lens was designed with an elliptic geometry to produce a wave focused to the center of the height dimension of the source and in the center of the chamber. The lens was 3D printed using stereolithography with a material known to allow for sufficient energy transmission from the element (PerFORM, Ceramic-Like Advanced HighTemp, Protolabs, Maple Plain, MN, USA) [36, 37]. The element was bonded to the lens using a thin epoxy layer. Wires were soldered to the inner and outer faces of the cylindrical element and connected to a BNC port. The sample holders were 3D printed using stereolithography (Clear Resin V4, Formlabs, Somerville, MA, USA). Small and large sample tubes had inner diameters of 6.35 mm and 9.525 mm, respectively, and both had a wall thickness of 1.59 mm (Tygon PVC E-1000, McMaster-Carr, Douglasville, GA, USA). Based on prior work, the tube materials were expected to have negligible pressure loss [25]. The tubes were sealed with stoppers on either end. The outer transducer housing was fabricated using selective laser sintering 3D printing (PA12 White, Protolabs).

### 2.3 Transducer Characterization and Pulse Generation

The electrical impedance of the transducer with the sample holder and sample tube in place with deionized degassed water in the exposure chamber was measured by an impedance analyzer (AIM4300, Array Solutions, Sunnyvale, TX, USA) connected by a coaxial cable. This measurement informed the design of an L-bridge matching network [35] for a matched electrical impedance Z = 10 Ω at a center frequency of 750 kHz. The transducer was driven by a multichannel class D amplifier controlled by an FPGA board (Cyclone DE1-SoC Board, Terasic, Hsinchu, Taiwan) delivering a unipolar square wave to the matching network. Custom MATLAB scripts were used to communicate with the FPGA and specify pulsing parameters.

A fiber-optic hydrophone (HF0690, Onda Corporation, Sunnyvale, CA, USA) was used to measure the acoustic pressure output of the transducer. Focal pressure waveforms were measured with deionized degassed water in the exposure chamber generating 40-cycle pulses with varying *p-*up to 11.6 MPa, beyond which the pressure could not be measured due to cavitation. For *p-*exceeding 11.6 MPa, the relationship between driving voltage and *p-*was quantified to linearly extrapolate *p-*up to the maximum amplifier output. It was determined that 15 V was needed to generate 1 MPa of pressure. An oscilloscope (TBS2000 series, Tektronix, Beaverton, OR, USA) was used to collect waveforms. The waveforms were filtered using a radiofrequency filter (BLP-1.9+, Lumped LC Low Pass Filter, DC - 1.9 MHz, 50Ω, Brooklyn, NY, USA). The waveforms were also averaged over 512 pulses to filter noise from the hydrophone, and the final waveform was reported. In the reported waveforms, the time axis was adjusted such that t = 0 represented the time of pulse delivery. Radial and axial 1D beam scans were performed along the major axes of the transducer to capture the focal beamwidths. The resulting beam profiles were recorded and normalized.

### 2.4 Cavitation Cloud Characterization

The cavitation cloud location and dimensions were determined using high-speed optical imaging. A high-speed camera (Nova S12 monochrome, Photron USA, San Diego, CA) with a 100 mm lens (Milvus 100 m f/1M ZF.2 Macro Lens, Zeiss, Jena, Germany) was used for imaging. A circular mirror was mounted to an optical rod at a 45° angle to the camera and the transducer’s axial dimension to capture images within the exposure chamber. The transducer was fixed to a clear frame that was positioned over a white LED strobe light (GS VitecMulti-LED QT Light, MultiLED G8 controller, 320 W power supply, Soden-Salmünster, Germany) for backlighting the images. The camera was mounted to an adjustable scissor jack fixed to an optics table for consistent alignment. 40-cycle pulses with *p-* = 21 MPa were delivered to the exposure chamber filled with 22 mL of deionized degassed water with a pulse repetition frequency (PRF) of 200 Hz. The camera was triggered once per pulse to collect images 2 μs after the end of the incident pulse reached the focus.

Images were analyzed to determine the location and size of the cavitation bubble cloud within the exposure chamber. Images were converted to binary first to identify and isolate the exposure chamber from the background. The exposure chamber was defined as a circular region with a centroid and diameter, and the surrounding image was masked and cropped outside of the diameter. The grayscale images of the cropped exposure chamber were then compared to a reference image captured before pulse delivery to remove any background artifacts and isolate the cavitation nuclei. The isolated images were then binarized, and the pixels with cavitation present were signified. This process was repeated over a series of images that were stacked to generate a heat map identifying the location of cavitation in the exposure chamber over 100 pulses.

### 2.5 FUSE Experimental Setup and DNA Extraction

Piscine and timber tissue were prepared for FUSE processing to validate the performance of the device for DNA extraction. For the piscine tissue study, fresh Atlantic salmon (*Salmo salar*) filets were prepared as they were previously [23]. Briefly, 25-50 mg cubes of tissue were sectioned and rinsed with deionized water. Samples were placed in a small sample tube (6.35 mm inner diameter) with lysis buffer composed of 270 μL of Buffer ATL and 30 μL Proteinase K (Qiagen Blood and Tissue Kit; Qiagen Incorporated, Hilden, Germany). For the timber tissue, live edge cuts of white oak (*Quercus alba*) were collected and prepared for FUSE processing two weeks after harvest. A file and rasp were used to collect shavings from the sapwood and cambium regions, as was done previously [26]. Shavings were passed through an ISO test sieve (Gilson Company Incorporated, Middleton, WI, USA) with square openings of 1 mm to separate any large segments from the shavings. 100 mg of shavings were placed in a large sample tube (9.525 mm inner diameter) with lysis buffer containing 1 mL of 1% PVP-40 Buffer AP1 solution and 8 μL of RNase A (Qiagen DNeasy Plant Kit; Qiagen Incorporated). Both sample tubes had stoppers on both ends to isolate the samples during FUSE processing.

The transducer exposure chamber was filled with 22 mL of deionized degassed water, and the sample tube assembly was suspended in the exposure chamber for processing using the sample holder aligned in the center of the chamber. 40-cycle pulses with *p-* = 21 MPa and a PRF of 200 Hz were delivered to the focus for sample processing and DNA release. Doses of 5,000 and 10,000 pulses were evaluated for both tissue types (n = 3), resulting in processing times of 25 and 50 seconds. After tissue processing, the sample holder was removed from the exposure chamber, and the tissue lysate was transferred to a 1.5 mL centrifuge tube for purification. Silica-column purification was done following the protocols recommended by the kit manufacturer (Qiagen Incorporated). The final elution volume was 200 µL for the piscine tissue and 80 µL for the timber.

### 2.6 Control DNA Extraction

Conventional DNA extraction methods were performed as a control to compare against FUSE using methods as was done in previous studies [26]. For piscine tissue samples, 25-50 mg of tissue was placed in a 1.5 mL centrifuge tube with 270 μL of Buffer ATL and 30 μL Proteinase K (Qiagen Blood and Tissue Kit; Qiagen Incorporated). Samples were incubated at 56 °C and vortexed every five minutes for 15 seconds until all tissue was disintegrated. This process took between 20-40 minutes. For timber tissue samples, 100 mg of shavings were processed using a mortar and pestle under liquid nitrogen. Liquid nitrogen was added to cool the mortar and pestle; then the tissue was placed in the mortar and homogenized for 30 seconds. The fractionated tissue was transferred to a 1.5 mL centrifuge tube with 1 mL of 1% PVP-40 Buffer AP1 solution and 8 μL of RNase A (Qiagen DNeasy Plant Kit; Qiagen Incorporated). Samples were incubated for one hour at 65 °C with a short vortex every ten minutes. Silica column purification was performed for both sample types (Qiagen Incorporated).

### 2.7 DNA Quantification

The released DNA was quantified by measuring the DNA yield and quality using the Qubit 4 Fluorometer (Thermo Fisher Scientific, Waltham, Massachusetts, USA) and NanoDrop One (Thermo Fisher Scientific). DNA yield was normalized based on the mass of sample input using values reported from the Qubit. DNA extract purity was evaluated by measuring the 260/280 and 260/230 ratios with the Nanodrop. An unpaired student’s t-test with unequal variance was used (p < 0.05) to evaluate the significance of the data collected with the Qubit and Nanodrop. Gel electrophoresis was also performed to assess the quality of the released DNA. For these experiments, 300 ng of DNA was used for piscine samples, while 1-11 ng of DNA was input for timber samples due to low DNA yields. The DNA was stained with 1x GelRed (Millipore Sigma, Burlington, Massachusetts, USA) and loaded into a 1% agarose gel in 1x TBE buffer (Thermo Fischer Scientific). The electrophoresis system was powered with 100 V for 1 hour. The ChemiDoc MP Imaging System (Bio-Rad, Hercules, CA, USA) was used to capture gel images, and the GeneRuler 1 kb Plus DNA Ladder (Thermo Fischer Scientific) was included to estimate the molecular weight of the input DNA.

### 2.8 Quantitative PCR Amplification

qPCR amplification was performed on DNA barcoding regions within the mitochondrial cyclooxygenase subunit 1 gene (COI) of *S. salar*, as well as the maturase K (matK) and ribulose bisphosphate carboxylase (rbcL) chloroplast genes of *Q. alba*. qPCR reactions were carried out as they were in prior studies for piscine and timber [26]. Briefly, in the piscine samples, the COI gene was targeted using a forward primer of 5′–CGCCCTAAGTCTCTTGATTCG–3′, and a reverse primer, 5′–GTAGTATGGTAATGCCTGCTGC–3′ that amplified a 536 bp region. Reactions contained 6 μL of 1X PowerUp SYBR Green Master Mix (Applied Biosystems, Thermo Fischer Scientific), 0.5 μM of each primer (Integrated DNA Technologies, Coralville, IA, USA), and 1.5 μL of template DNA. The thermal cycler was programmed as follows: 50 °C for 2 minutes, 95 °C for 2 minutes, and 30 cycles of 95 °C for 15 seconds and 60 °C for 1 minute. Amplifications with threshold cycle (Ct) values less than or equal to 25 cycles were considered successful.

For timber samples, the matK gene was amplified using a forward primer, 5’-TTTCCGGTCATCCCATGCTTT-3’, and reverse primer, 5’-TGCAGGATTTCGTCGAACACT-3’, that targeted a 243 bp region. For rbcL targeting, a forward primer, 5’-ACGATGCTACCACATCGAGC-3’, and reverse primer, 5’-GAGGCGGACCTTGGAAAGTT-3’, were used to amplify a 212 bp region. qPCR mixture ratios followed those of the piscine reactions, but for this case a total volume of 40 μL was used for all reactions due to lower template DNA quantities. For both matK and rbcL primers, thermal cycling was performed using the following parameters: 50 °C for 2 minutes, 95 °C for 2 minutes, and 45 cycles of 95 °C for 15 seconds, 57 °C for 15 seconds, and 72 °C for 1 minute. Samples with Ct values less than or equal to 40 cycles were considered successful. All amplifications for both timber and piscine samples were performed in triplicate, and each qPCR run included negative controls containing nuclease-free water instead of template DNA.

## 3. Results

### 3.1 Transducer Output Characteristics

Electroacoustic and acoustic propagation simulations suggest that the transducer design can generate sufficient focal gain for cavitation generation. Electroacoustic simulations estimated a surface pressure of 0.2 MPa, at a driving voltage of 200 V. Acoustic propagation simulations demonstrated the spatial and temporal evolution of the pressure field throughout the cylindrical exposure chamber. The simulated pressure waveforms exhibited interference patterns during the delivery of a multi-cycle pulse, and beam profiles show that the acoustic focus is expected to be contained within the sample tube (**Figure 2**). The simulated pressure waveform demonstrates that *p-*occurred approximately 47 μs after pulse delivery. At *p-*, the simulated focal gain was 60, consistent with other focused ultrasound devices [32, 38, 39]. The simulated beam profiles reveal that the focus is expected to be centered along the radial and axial dimensions of the chamber. Results show that beam plots had -6 dB beam dimensions of 11.6 mm and 0.75 mm along the axial and radial dimensions of the transducer.

**Figure 2.**
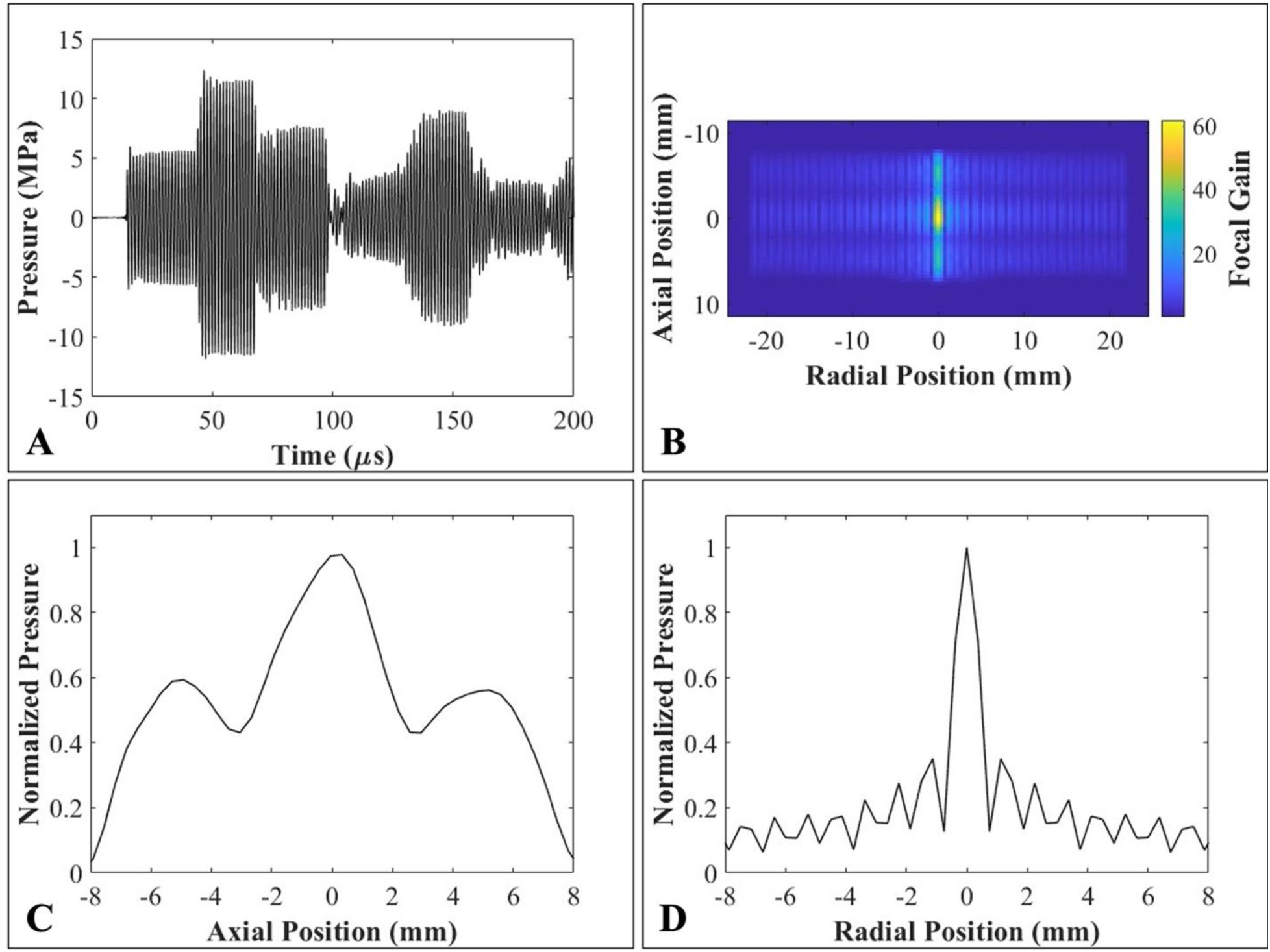
Simulated waveform and focal pressure output. (A) Pressure waveform in the geometric focus of the transducer assuming *p_0_* = 0.2 MPa. (B) Cross-sectional focal gain in the geometric focus. (C) Axial and (D) radial beam profiles show that the focus is expected to be in the center of the transducer with -6 dB beam dimension of 11.6 mm axially and 0.75 mm radially. The plotted focal gain is the ratio of simulated focal pressure to source surface pressure.

The fabricated transducer was cylindrical with a focal length of 19.25 mm. Consistent with the simulated output, measured pressure waveforms demonstrated that 750 kHz multi-cycle pulses produced interference patterns with variable pressure amplitudes throughout the pulse duration (**Figure 3A**). The beam profiles showed -6 dB beamwidths of 1.2 mm and 4.8 mm in the radial and axial dimensions, respectively (**Figure 3B-C**). Measured beamwidths were used to predict the bubble cloud dimensions [30, 31], and these results suggest that cavitation should be confined within the sample tube. The measured beam profiles were approximately 60% different than simulations. It is expected that differences between the measured and simulated beamwidths are due to slight asymmetry in the fabricated transducer and simplifications in the model, which may affect the complex patterns of wave propagation and interference in the exposure chamber. The measured beam profiles also showed that the acoustic focus was centered in the radial dimension but shifted toward the bottom of the chamber in the axial dimension.

**Figure 3.**
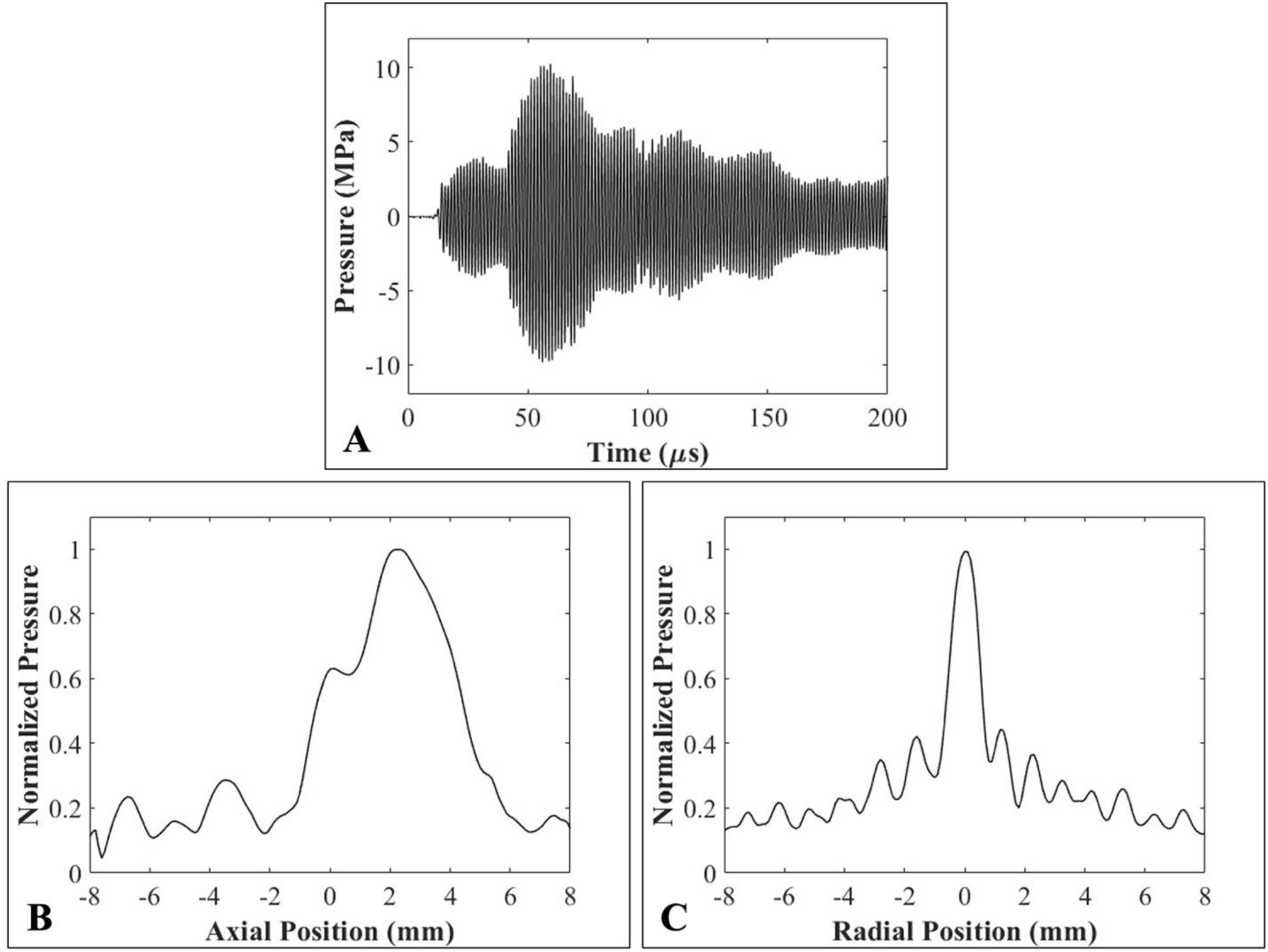
Measured pressure output. (A) Pressure waveform in the geometric focus of the transducer with *p-* = 9.9 MPa. (B) Axial and (C) radial beam profiles with -6 dB dimensions of 4.8 mm axially and 1.2 mm radially.

### 3.2 Cavitation Cloud Characteristics

High-speed optical imaging experiments demonstrated that the fabricated FUSE device generated consistent cavitation (**Figure 4**). Imaging was done axially to characterize the cavitation cloud in the radial and angular dimensions of the cylindrical transducer. In agreement with the simulated and measured beam profiles, cavitation was observed in the center of the chamber. Over 100 pulses, cavitation was observed 100% of the time. The cavitation cloud was dynamic and characteristic of cavitation-based focused ultrasound techniques [40]. The mapped spatial distribution of cavitation events showed the size and location of the cavitation cloud within the exposure chamber. Results show that all cavitation events were contained within the bounds of the tube, demonstrating that the device met the design requirement of generating cavitation within a small sample tube.

**Figure 4.**
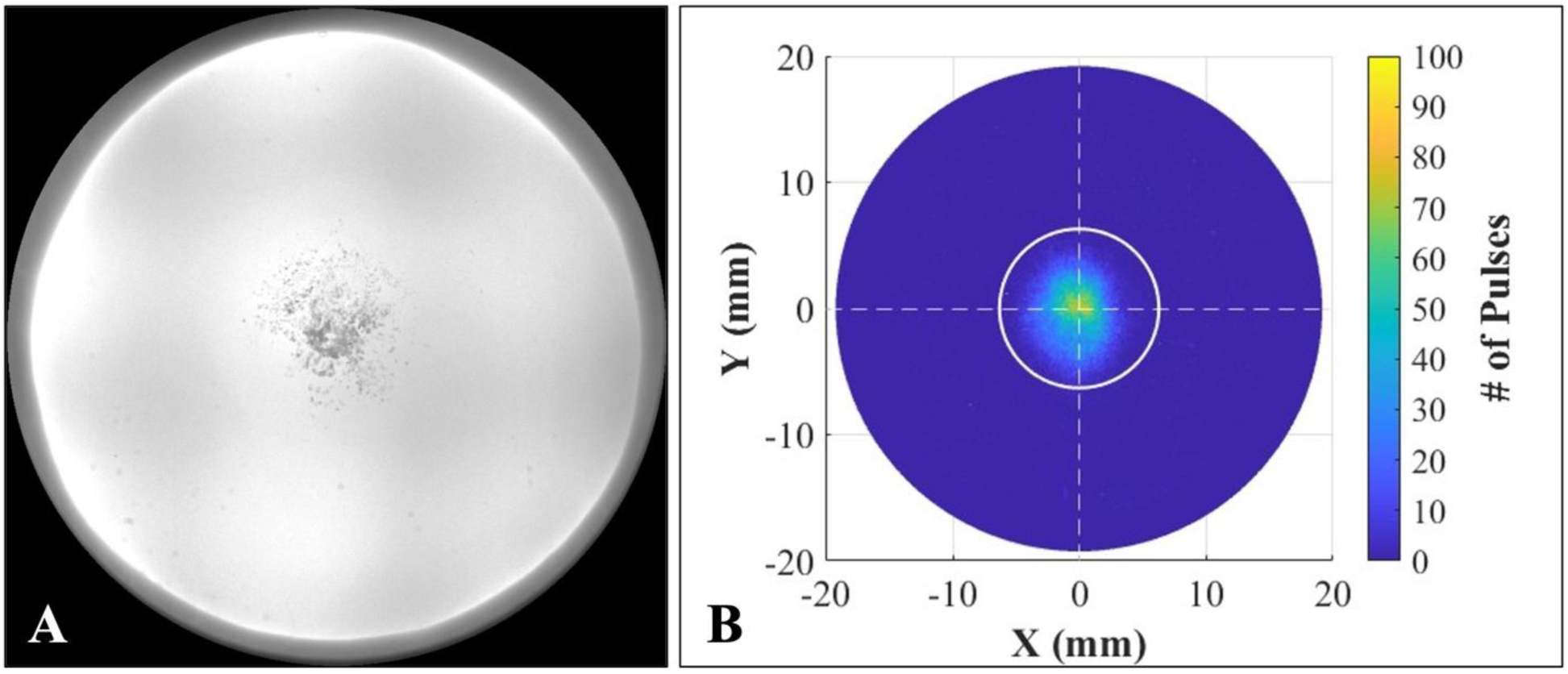
Cavitation bubble cloud imaging. (A) High-speed optical imaging was performed to capture the cavitation bubble cloud 2 μs after the completion of a 21 MPa, 40-cycle pulse to the acoustic focus. (B) The location of cavitation events was mapped over 100 pulses to show the size and position of the cavitation cloud within the exposure chamber. The white circle indicates the size of a small sample tube with an inner diameter of 6.3 mm. Results show that cavitation is sustained within the sample tube.

### 3.3 DNA Extraction and Feasibility of PCR-based Detection

The feasibility of this device for releasing DNA from piscine and timber tissue was evaluated to demonstrate the potential of FUSE DNA sample preparation in a miniaturized system (**Figure 5**). For both tissue types, tissue disruption was observed after 25 and 50 seconds of FUSE processing. For piscine tissue, complete disintegration of all visible tissue fragments was achieved after 25 seconds of processing. For timber tissue, the extent of tissue breakdown could not be determined visually. However, tissue circulation was observed during treatment, consistent with prior work demonstrating that tissue circulation is indicative of cavitation activity and tissue breakdown [26]. Tissue homogenization was evidenced by the resulting DNA yield from both tissue types (**Figure 6**). For the piscine samples, high quantities of DNA were released after 25 and 50 seconds of processing, with FUSE DNA yields comparable to control methods. DNA yields from timber samples were also comparable between FUSE and control methods. However, increasing the FUSE processing time from 25 to 50 seconds significantly increased DNA yield from timber samples (p < 0.05). Due to high variability in the control DNA yield results for timber, there were no significant differences between the controls and either of the FUSE groups.

**Figure 5.**
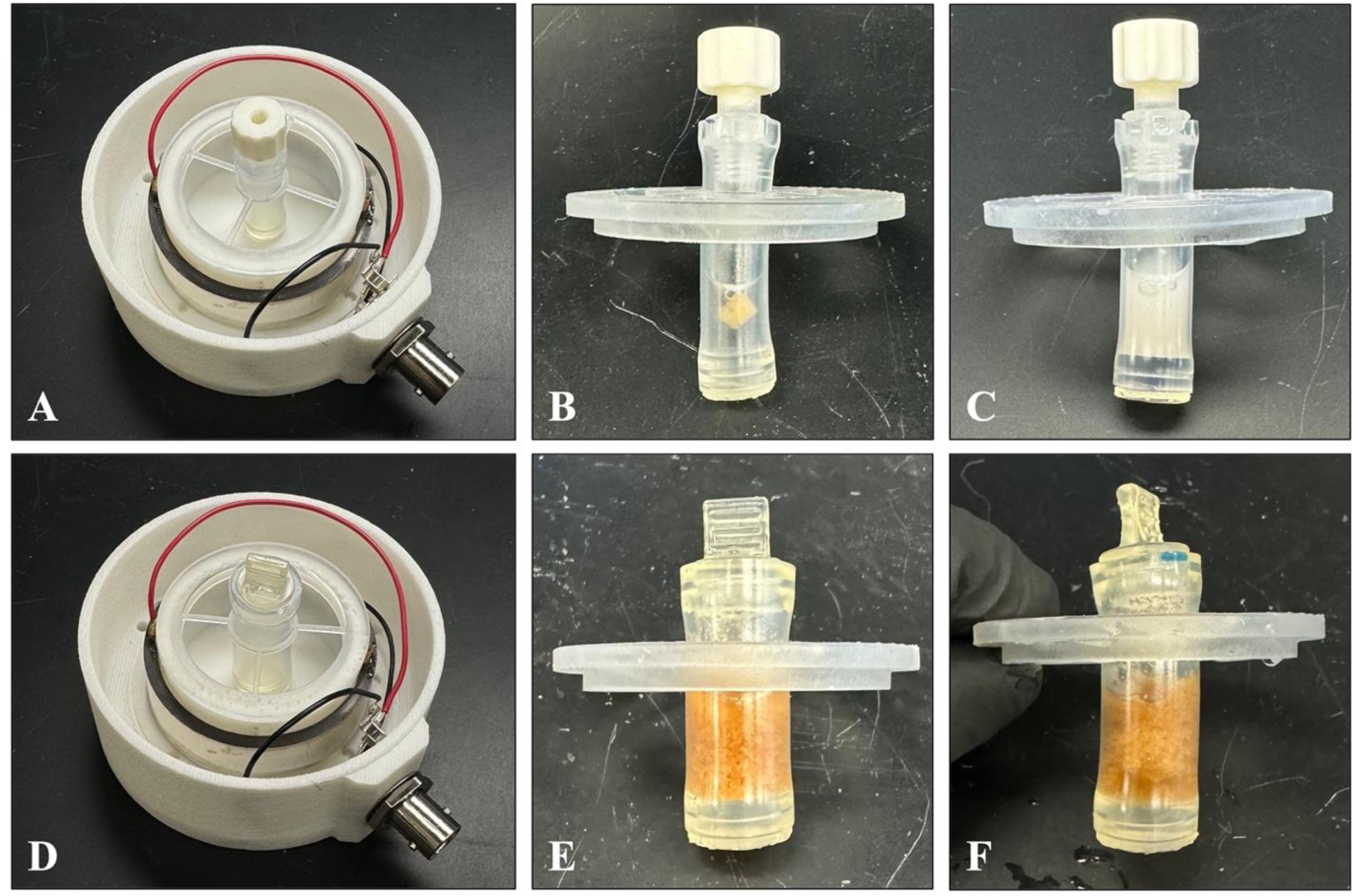
FUSE sample preparation and processing. The FUSE experimental configuration for the preparation of piscine (A-C) and timber (D-F) tissue. In the first column the device is shown with a (A) 6.35 mm tube and (D) 9.525 mm tube setup. The second column shows the (B) piscine and (E) timber tissue prepared in lysis buffer. The third column demonstrates the (C) piscine and (F) timber tissue after FUSE tissue processing.

**Figure 6.**
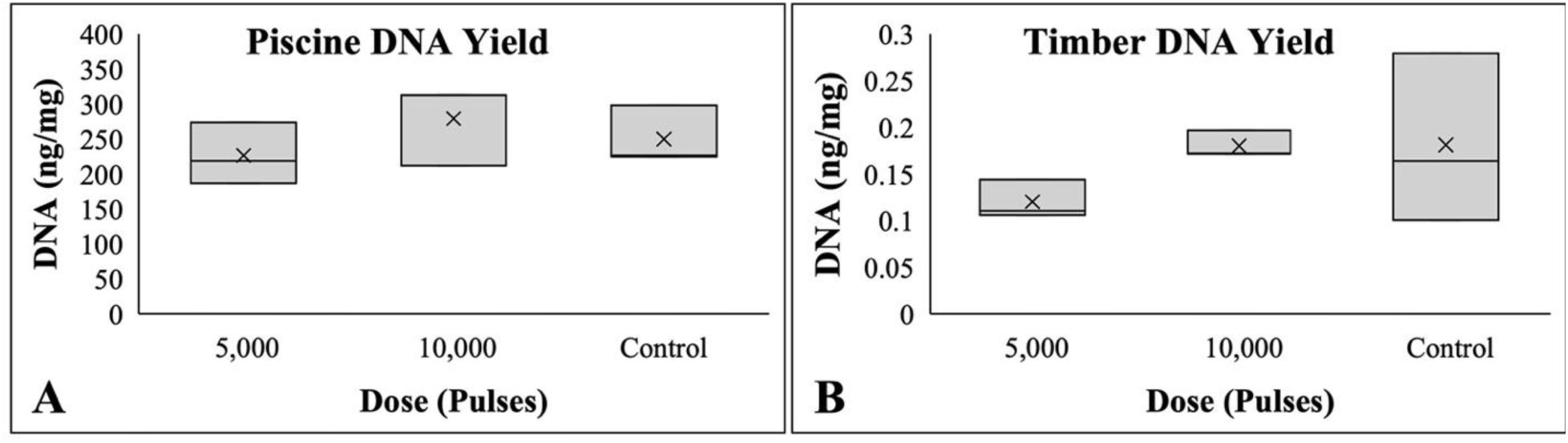
Piscine and timber DNA yield. (A) Piscine and (B) timber DNA yield results demonstrate that the miniaturized FUSE device released DNA from both tissue types with yields comparable to conventional extraction methods, represented by the controls. For the timber samples, 10,000 pulses released significantly more DNA than 5,000 pulses (p < 0.05).

DNA purity results were also reported for piscine and timber samples (**Table 1**). 260/280 and 260/230 ratios were within the expected norms for the piscine samples. In contrast, for the timber samples, 260/280 and 260/230 ratios were lower than the anticipated values for DNA for both FUSE and control groups. No significant differences in 260/280 ratios were observed between treatment groups for either sample type. However, timber samples processed with FUSE for 25 seconds exhibited significantly higher 260/230 ratios than those processed for 50 seconds or prepared using control methods.

**Table 1.**
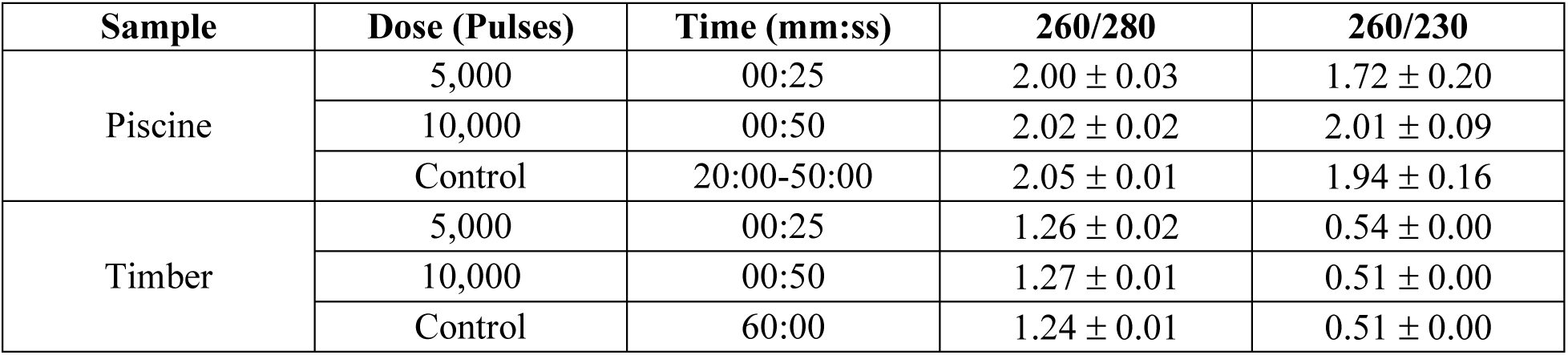
Piscine and timber DNA purity ratios. 260/280 and 260/230 ratios were reported for piscine and salmon samples prepared with FUSE and controls. Results show that DNA purity ratios are within the expected range for piscine samples, while timber ratios are lower than what is expected for DNA for both FUSE and controls.

Gel electrophoresis was also performed to assess DNA quality. This analysis was limited to piscine samples, as DNA yields from timber samples were insufficient for gel electrophoresis.

Results show that increasing processing time fragmented the DNA (**Figure 7**). However, when samples were prepared using 25 seconds of processing, high molecular weight bands were still present with sizes as great as 20 kB.

**Figure 7.**
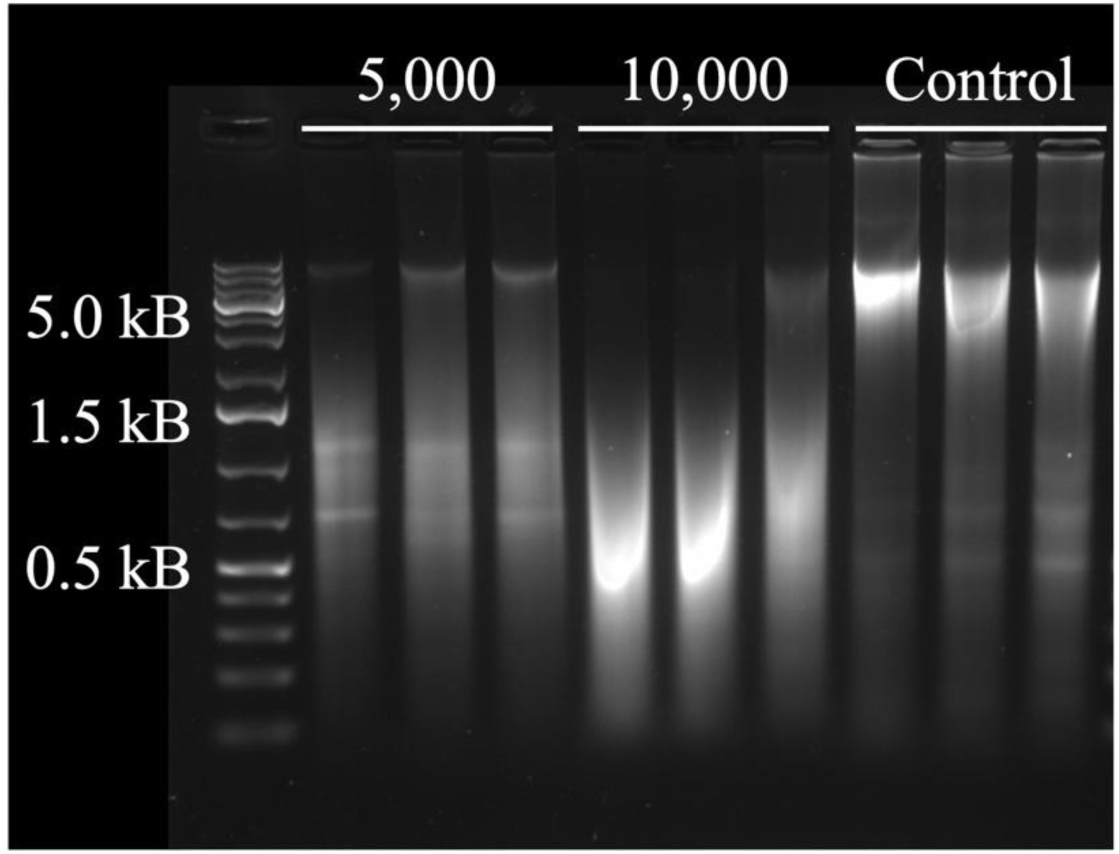
Piscine gel electrophoresis. Gel electrophoresis results show that increasing the FUSE processing time from 5,000 to 10,000 pulses resulted in DNA shearing. The effect of 5,000 pulses on DNA shearing is comparable to controls.

Successful qPCR amplification was achieved for both piscine and timber samples, demonstrating the potential of the fabricated FUSE device to prepare DNA suitable for downstream detection assays (**Table 2**, **Table 3**). Both 25- and 50-second FUSE treatments resulted in successful qPCR amplification for both sample types. For piscine samples, results suggest that increasing the dose improved qPCR success rates, and the amplification efficiency across treatment groups was comparable. For the timber samples, a qPCR success rate of 100% was observed for samples across both FUSE treatment groups. In contrast, the control group had an overall qPCR success rate of 39%. Further, qPCR amplification efficiency was also significantly higher for FUSE processed samples than for controls using both the matK and rbcL primer sets. For timber samples processed with FUSE for 25 seconds, amplification efficiencies were 25.0 ± 0.34 and 24.6 ± 0.15 for the matK and rbcL genes, respectively, compared with 39.2 and 33.0 ± 2.12 for samples prepared using control methods.

**Table 2.**
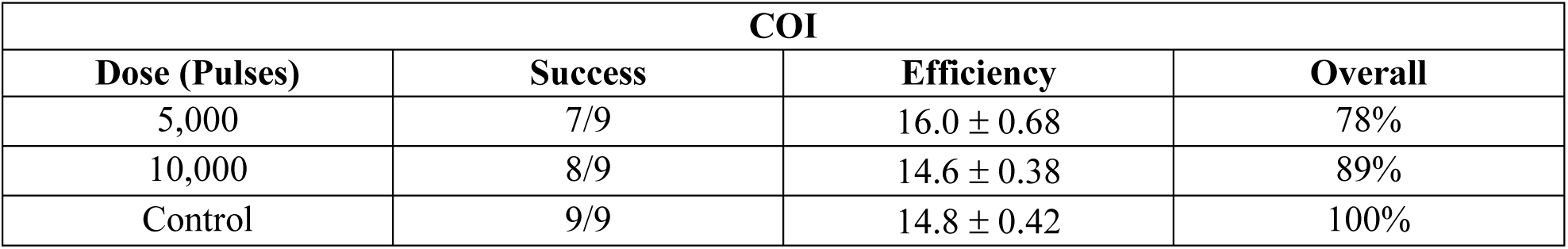
qPCR amplification of the COI gene in piscine samples. The qPCR success rate and efficiency in amplifying the COI mitochondrial gene is shown. Results show that qPCR success and efficiency increased with increasing FUSE processing time.

**Table 3.** qPCR amplification of matK and rbcL genes in timber samples. The qPCR success rate and efficiency in amplifying the matK and rbcL chloroplast genes is shown. Results show that qPCR was 100% successful in both genes after FUSE DNA release of timber samples. qPCR efficiency was not affected by FUSE processing time.

|  | matK |  | rbcL |  |  |
| --- | --- | --- | --- | --- | --- |
| Dose (Pulses) | Success | Efficiency | Success | Efficiency | Overall |
| 5,000 | 9/9 | $25.0 \pm 0.34$ | 9/9 | $24.6 \pm 0.15$ | 100% |
| 10,000 | 9/9 | $26.7 \pm 0.70$ | 9/9 | $24.9 \pm 0.22$ | 100% |
| Control | 1/9 | 39.2 | 6/9 | $33.0 \pm 2.12$ | 39% |

## 4. Discussion

This study presented the design of a portable transducer developed for the preparation of DNA from complex tissues in POC settings to broaden the accessibility and applicability of NAATs. The device was designed using a cylindrical transducer element capable of delivering 750 kHz pulses with a pulse duration of 40 cycles to an enclosed sample tube. The overall size of the device was small (77 x 31 mm) to allow for easy transport and versatile use. Characterization of the acoustic output showed radial and axial beamwidths of 1.2 mm and 4.8 mm, respectively. Cavitation was characterized using high-speed optical imaging, and results showed that the bubble cloud was generated in the center of the sample chamber and confined within the bounds of the sample tube dimensions. The performance of the device was validated by demonstrating the feasibility of FUSE for piscine and timber tissue processing, DNA release, and qPCR amplification. For both sample types, DNA with yields, purity, and quality sufficient for qPCR amplification was achieved after only 25 seconds of FUSE processing. For timber samples, FUSE significantly improved PCR performance compared with controls, suggesting improved DNA quality. Overall, these results demonstrate that the portable FUSE device can extract DNA suitable for PCR from piscine and timber samples.

The transducer design used in this study was selected primarily based on utility and ease of use as well as the size and shape of the desired sample tube. In prior FUSE studies, a large 32-element hemispherical transducer was used to deliver focused ultrasound pulses to a sample of interest, and the sample tube was aligned in the focus such that the length of the tube was perpendicular to the face of the transducer [23–26]. This configuration required a robotic positioning system to expose the entire sample volume to the focus. To prevent the need for movements during tissue processing, the miniaturized transducer was designed such that the major axis of the bubble cloud spanned the height of the sample tube. This was done with a cylindrical source to increase the focusing strength and the size and density of the cavitation bubble cloud [30]. Previous FUSE studies employed a transducer with an f-number of 0.62, whereas the transducer developed in this study had an f-number of 0.5, providing stronger geometric focusing. Comparison of the simulated acoustic output and measured pressure waveforms revealed discrepancies between the modeled and measured beam profiles. These differences are expected to be due to simulation design simplifications and experimental variability. One limitation of the simulation was that it did not explicitly account for the lens material on the inner surface of the cylindrical piezoelectric source. Instead, the source was modeled with an elliptical geometry and an arc centered along the height dimension to approximate the focusing effects of the lens. During device fabrication, slight misalignment between the piezoelectric element and the elliptical arc of the lens may have contributed to asymmetric focusing. Experimental variability, including differences in water height, dissolved gas concentration, and water temperature, may have further contributed to discrepancies between the simulated and measured results. Future work intends to improve the model such that the source element is flat and positioned along a curved lens. The relative positioning of the source element and lens curvature will also be evaluated to more accurately represent acoustic wave propagation within the device.

The cylindrical geometry of the transducer and exposure chamber also influenced the acoustic pressure field through wave reflections and interference patterns. It is expected that the applied pulsing schemes and acoustic properties of the medium in the exposure chamber contributed to interference between incident and reflected waves at the focus. Future work will investigate alternative pulsing schemes to maximize constructive interference and increase focal pressure. Leveraging these interference patterns may provide a means of increasing focal pressure without excessive power and voltage requirements, an important consideration for the development of a portable POC device.

The DNA extraction results confirmed that a portable FUSE device effectively and rapidly releases DNA from various complex sample types. The successful DNA extraction from timber tissue in just 25 seconds was particularly notable given the challenges associated with processing timber samples. Previous FUSE studies required five minutes of processing at a PRF of 1 kHz, corresponding to 300,000 pulses, to release DNA from timber [26]. In contrast, the present study required 25 seconds and only 5,000 pulses delivered at 200 Hz, a 60X reduction in the number of pulses. The timber qPCR results further validated the performance of this device in preparing DNA from robust tissues. Compared with control methods, FUSE processing resulted in more consistent amplification and lower Ct values, suggesting that the extracted DNA was more suitable for downstream detection. These findings are particularly promising given the short processing times required for FUSE. Ongoing work will include additional experimental replicates to confirm the reproducibility of this trend.

The piscine DNA extraction results demonstrated that FUSE can preserve high-molecular-weight DNA under certain processing conditions. Previous studies investigating FUSE have reported DNA fragmentation following acoustic processing [24–26]. In contrast, gel electrophoresis in the present study revealed minimal DNA shearing after 5,000 pulses, with high-molecular-weight DNA bands observed. These findings suggest that the extent of DNA fragmentation may be influenced by the applied acoustic parameters. Future work will further characterize the relationship between FUSE processing parameters and DNA integrity.

Overall, the transducer developed in this study shows great promise for the development of a portable FUSE platform capable of enabling rapid DNA release from complex sample types. This FUSE device has the potential to broaden the accessibility and versatility of NAATs for several applications including environmental security and global health. A rapid and reliable DNA extraction method capable of preparing robust sample types, field performance (battery-powered), and operation by non-technical users is necessary to improve the current capabilities of NAATs [8]. The device presented in this study addresses many of the key factors necessary to enable POC use including small size, extraction in less than 30 seconds, and the ability to prepare complex samples for reliable and efficient qPCR assays. Ongoing work aims to further refine this device to enhance portability and field performance. Future studies will focus on identifying the acoustic parameters that enable optimal sample preparation while limiting power requirements to allow for the development of a battery-powered amplifier for using the device in low-resource settings.

## 5. Conclusion

A miniaturized, portable device for FUSE DNA sample preparation was designed and fabricated to address the limitations of NAATs. The transducer pressure output and beam profiles were measured with focal hydrophone measurements. High-speed optical imaging was used to validate that the focal gain was great enough to generate sustained cavitation, and results show that cavitation was confined within the planned dimensions. FUSE DNA extraction with piscine and timber samples demonstrated that this device can release high quantities of high-quality DNA suitable for PCR amplification in 25 seconds. These results demonstrate the potential of this device as a novel POC DNA extraction platform capable of preparing complex sample types, warranting the continued development of FUSE for new indications and enhanced performance.

## 6. Acknowledgements

Funding for this work was provided by the Gordon and Betty Moore Foundation (grant #8518). The authors would like to particularly thank the Moore Foundation’s Science Program for their ongoing support of this project. We would also like to acknowledge Conservation X Labs, the Virginia Tech Department of Biomedical Engineering and Mechanics, and the Virginia Tech Institute for Critical Technology and Applied Science for their support of this work. We would also like to acknowledge Truman Lumber for providing the timber samples used in this study.

